# A negative feedback loop between Insulin-like Growth Factor signaling and the lncRNA SNHG7 tightly regulates transcript levels and proliferation

**DOI:** 10.1101/709352

**Authors:** David N. Boone, Andrew Warburton, Sreeroopa Som, Adrian V. Lee

**Affiliations:** Women’s Cancer Research Center; Department of Biomedical Informatics, University of Pittsburgh; University of Pittsburgh Cancer Institute Academy/Computer Science, Biology, and Biomedical Informatics Academy; Icahn School of Medicine at Mount Sinai; Stanford University; Department of Pharmacology and Chemical Biology, University of Pittsburgh; University of Pittsburgh Cancer Institute

**Keywords:** Long non-coding RNAs, Insulin-like Growth Factor, Breast Cancer, Proliferation, SNHG7

## Abstract

Evidence suggests Insulin-like growth factor 1 (IGF1) signaling is involved in the initiation and progression of a subset of breast cancers by inducing cell proliferation and survival(1, 2). Although the signaling cascade following IGF1 receptor activation is well-studied(3, 4), the key elements of the transcriptional response governing IGF1’s actions are not well understood. Recent studies reveal that the majority of the genome is transcribed and that there are more long non-coding RNAs (lncRNAs) than protein coding genes(5), several of which are dysegulated in human cancer(6, 7). However, studies on the regulation and mechanism of action of these lncRNAs are in their infancy. Here we show that IGF1 alters the expression levels of a subset of lncRNAs. SNHG7, a member of the small nucleolar host gene family, is a highly-expressed lncRNA that is consistently and significantly down-regulated by IGF1 signaling by a post-transcriptional mechanism through the MAPK pathway. SNHG7 regulates proliferation of breast cancer cell lines in a dose-dependent manner, and silencing SNHG7 expression causes cell cycle arrest in G0/G1. Intriguingly, SNHG7 alters the expression of many IGF1 signaling intermediates and IGF1-regulated genes suggesting a feedback mechanism to tightly regulate the IGF1 response. Finally, we show with TCGA data that SNHG7 is overexpressed in tumors of a subset of breast cancer patients and that these patients have lower disease-free survival than patients without elevated SNHG7 expression. We propose that SNHG7 is a lncRNA oncogene that is controlled by growth factor signaling in a feedback mechanism to prevent hyperproliferation, and that this regulation can be lost in the development or progression of breast cancer.

**SIGNIFICANCE STATEMENT:** IGF1 signaling drives proliferation and survival and is important for the initiation and development of a subset of breast cancers. IGF1 is known to control the expression of thousands of protein coding genes, but it is unknown if it alters the expression of other gene types, such as long noncoding RNAs. Here we demonstrate that IGF regulates lncRNAs including the mostly unstudied SNHG7. We further show that SNHG7 is necessary for proliferation and modulates IGF1 signaling through a novel feedback mechanism that is required for fine-tuning of the transcriptional response to growth factor signaling and proliferation of breast cancer cells. SNHG7 is highly expressed in a subset of breast cancer patients with poor prognosis giving further credence that it is a novel oncogene.

## INTRODUCTION

Substantial evidence implicates IGF1 signaling in the initiation and development of a number of cancers including breast cancer (4). The signaling initiated by IGF1 binding to IGF1R, a receptor tyrosine kinase, is well known. IGF1R activation induces a phosphorylation cascade through IRS1 and IRS2, which stimulates the MAPK and PI3K/AKT pathways(3). Ultimately, IGF1 signaling leads to a robust and temporal transcriptional response(8, 9)—10% of all protein coding genes(9)—and an array of biological processes including cell proliferation and survival(10). While the signaling and biological responses elicited by IGF are well-known, the IGF-regulated genes and the molecular mechanisms that govern those biological responses are largely unclear. Furthermore, there has not been a comprehensive examination of IGF1-induced transcriptome changes using RNA sequencing. This is critical given that IGF regulates a vast number of protein coding genes and recent large-scale omics studies including ENCODE demonstrate that there are more non-coding transcripts than coding(5, 11, 12).

Long non-coding RNAs (lncRNAs) are a diverse class of RNA molecules that are loosely defined by an arbitrary length of greater than 200 nucleotides and the apparent lack of protein coding potential(13–17). The number of lncRNAs, although debated in the literature, at least rivals the number of protein coding genes. While the vast majority were recently identified and do not have a known function, several lncRNAs including XIST(18–20), HOTAIR(7), and H19(21, 22), have been studied for decades. From those and recent studies, it is evident that lncRNAs are important regulators of a variety of cellular processes including transcriptional regulation, chromatin structure, RNA stability, and cell proliferation through a variety of novel mechanisms that often are due to the ability of lncRNAs to bind to DNA, RNA, and proteins and act as guides, scaffolds, and decoys(23). Further, the dysregulation of lncRNAs is implicated in the development and progression of many diseases including breast cancer(6, 7, 13, 24–29). Therefore, it is imperative to identify and characterize the regulation and functional significance of novel lncRNAs to understand basic biological processes and the pathogenesis and treatment of diseases such as breast cancer.

There has not been a comprehensive examination of regulation of lncRNAs by IGF1, but IGF/Insulin signaling represses the expression of CRNDE(30), a lncRNA highly expressed in colorectal cancer and gliomas(31, 32). In this report we aimed to further understand the molecular mechanisms of the biological functions of IGF1 and to leverage the extensive knowledge of IGF1 as a model system to identify and characterize growth factor regulated lncRNAs that are functionally critical lncRNAs in breast cancer. Here, we demonstrate through whole transcriptome RNAseq that IGF1 signaling regulates a subset of lncRNAs that are altered in breast cancer. Further, we show that the known but unstudied lncRNA, SNHG7, which is amplified or overerxpressed in ~5% of breast tumors in TCGA, is downregulated by IGF through a post-transcriptional mechanism through MAPK and controls proliferation in a dose-dependent manner. SNHG7, in part, tightly controls proliferation by altering mRNA levels of both IGF1 signaling intermediates and downstream IGF1 regulated genes. Thereby, we identified a novel fine-tuning feedback mechanism of growth factor induced proliferation and gene expression response that is disrupted in the tumors of a subset of breast cancer patients.

## RESULTS

### IGF regulates lncRNAs that are dysregulated in breast cancer

The MCF7 cell line is a model breast cancer cell line that is robustly responsive to IGF1. Addition of IGF1 to serum deprived MCF7 cells leads to rapid activation of AKT/PI3K and MAPK pathways, expression changes of 1000s of genes, and proliferation (9). To identify lncRNAs regulated by IGF1 signaling that may be critical for proliferation of breast cancer cells, we examined the transcriptional response induced by the addition of IGF1 to serum starved MCF7 cells after 3 and 8hrs using whole transcriptome RNAseq. The Tuxedo package(33) was used for transcriptome assembly and differential gene expression analysis. The reads were aligned and transcriptomes assembled using the GRCh38 genome build with all annotated Gencode v21(34) transcripts allowing for novel transcript detection. Additionally, reads that mapped to tRNAs, snoRNAs, miRNAs, and rRNAs were masked during transcript assembly to ensure proper expression calls of lncRNAs that are ‘hosts’ for small noncoding RNAs (Fig. 1A). When small ncRNA reads were not removed, expression of host lncRNA genes were often miscalculated because of the abundant reads of the small ncRNAs that are present in their introns, but are not part of the mature lncRNA (data not shown). IGF1 signaling significantly (q<0.05) induced a greater than 1.5-fold change in 1067 and 2061 annotated (Gencode v.21) genes at 3 and 8 hrs respectively (Fig. S1A-B; Supplementary Table 1). Individual gene expression changes were validated by qPCR in the same, and in an independent set of RNA (Fig. S1C). The global changes in gene expression observed correlated with the changes shown by expression microarray in our previous study(9) (data not shown). Also, as expected, pathway analysis of IGF1-regulated genes at 3 and 8hrs (FDR<0.05;FC>2.0) revealed that these transcripts were involved in activation of proliferation, survival, and cancer development, as well as, inhibition of cell death (Fig. S1D). Collectively, the qPCR and pathway analyses demonstrate the quality and validity of the RNA-seq data.

**Figure 1.**
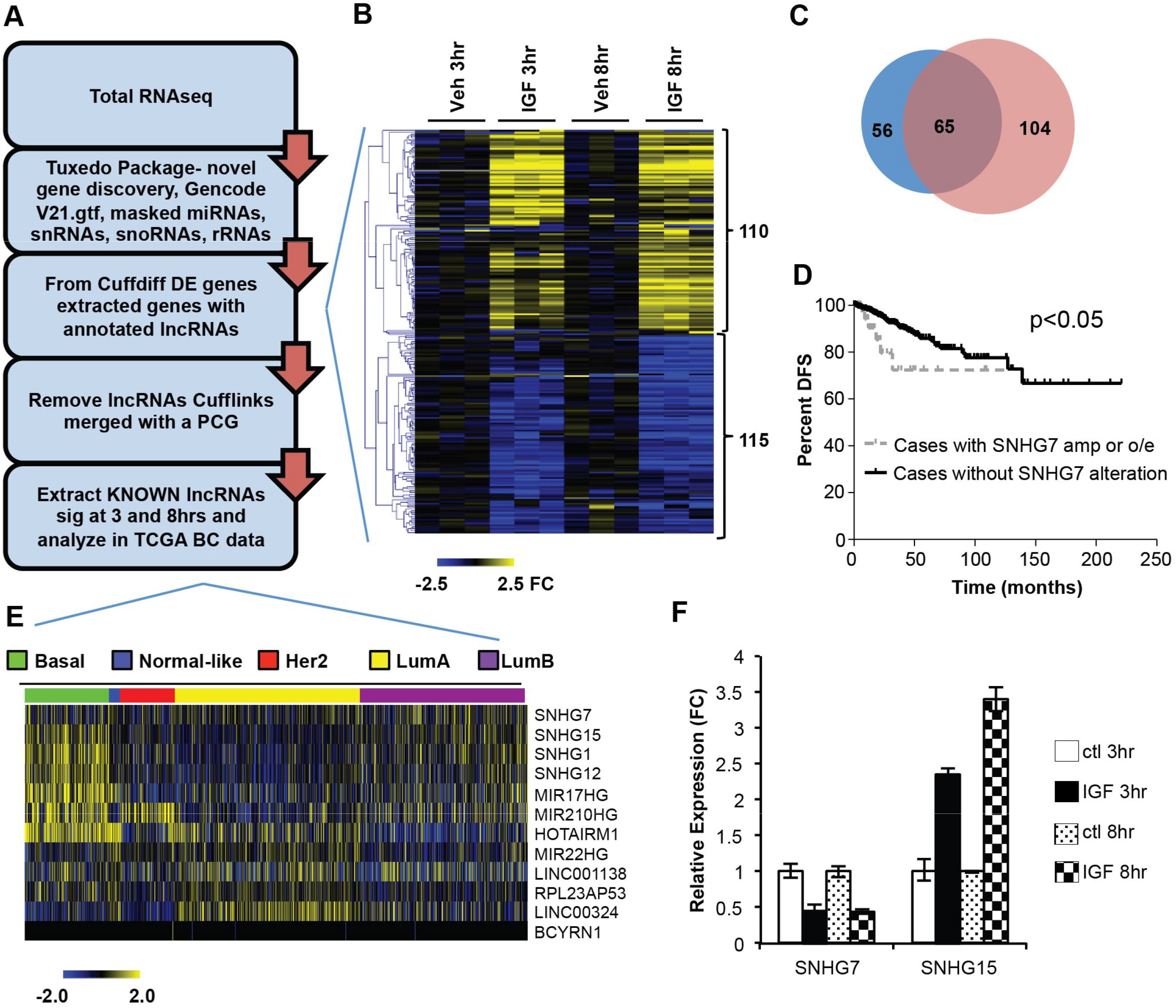
IGF1 Signaling regulates the expression of lncRNAs. (A) RNAseq and informatics pipeline used to identify persistently IGF1 regulated known lncRNAs. The Tuxedo package was used to determine differentially expressed (DE) genes after IGF1 treatment. Novel gene discovery was allowed, but for a conservative estimate only genes with Gencode V21 lncRNA annotation that did not overlap with a protein coding gene (PCG) annotation on either strand are reported. (B) Heatmap of the expression of lncRNAs (as defined in Fig. 1A) significantly regulated by IGF1 treatment at 3 or 8hrs. Expression levels are normalized to the mean of the respective vehicle (Veh) control. Each column is a replicate of the indicated treatment group and each row is an individual lncRNA (C) Venn Diagram demonstrating the number of lncRNAs significantly regulated at 3hrs (blue), 8hrs (red), or both (purple). (D) SNHG7 is amplified or overexpressed in a subset of the tumors of TCGA patients (N=45). Those patients have a worse Disease Free Survival (Log-rank Test p<0.05) than patients with normal levels of SNHG7 DNA and RNA (N=866). Patients with a copy number loss of SNHG7 (N=3) were ignored. (E) Normalized RNASeq V2 RSEM expression data from annotated lncRNAs in TCGA breast cancer (BC) data that are regulated by IGF at 3 and 8hrs was downloaded from the TCGA data portal. Values were log2 transformed and then median centered by gene. Breast cancer molecular subtypes determined by PAM50 scores(47) are indicated by color. (F) Validation of IGF regulation of indicated lncRNA by qPCR. Results are reported as the mean expression normalized to time-matched vehicle control +/- SD (ttest p<0.05 for all comparisons to respective control).

To determine if any of the differentially expressed genes were lncRNAs, we used a conservative approach of extracting any IGF-regulated gene that was annotated as a lncRNA in Gencode v.21 that was not merged with a protein coding gene during transcript assembly (Fig. 1A) thus excluding many highly-overlapping antisense lncRNAs that were not properly aligned due to the use of an unstranded RNAseq library. This revealed that the expressions of 225 previously annotated lncRNAs with a minimum fpkm of 1 at either 3 or 8hrs were significantly altered by IGF1 treatment at 3 or 8hrs with nearly an equal number upregulated as downregulated (Fig. 1B; Supplementary Table 2). Consistent with mRNA regulation by IGF in this and our previous study, slightly more were significantly regulated at 8hrs than 3hrs (Fig. 1C). The expression of 65 annotated lncRNAs changed at both 3 and 8hrs suggesting early and sustained control by IGF1 signaling (Fig. 1C). To identify cancer relevant, IGF-induced lncRNAs, we sought to examine the alteration of these lncRNAs in The Cancer Genome Atlas (TCGA) breast cancer data (http://cancergenome.nih.gov/). Of the 65 lncRNAs only 12 had a “KNOWN” gene status by GENCODE meaning the annotation is identical to a known and curated gene in Entrez and is reported in TCGA. Examination of the 12 lncRNAs in the TCGA breast cancer data through the cBIO portal(35, 36) revealed that 11 of them have copy number or gene expression alterations in a subset of breast cancer patients (Table 1). Interestingly, the dysregulation of one lncRNA, SNHG7, is enriched in a patient population with a poorer prognosis. SNHG7 is altered in ~5% of all breast cancer tumors in TCGA (70 of 1105 samples; 67 overexpressed or amplified). Patients with overexpressed or amplified SNHG7 had a statistically significant poorer disease-free survival (Fig. 1D and Table 1; logrank test p-value=0.0139; N=7 of 45 with altered SNHG7 relapsed vs. 61 of 866). This demonstrates that SNHG7 is potentially translationally relevant and was selected for further study. In addition, analysis of gene expression data extracted for all TCGA breast cancer samples demonstrates that the expression of many of the 12 IGF-regulated lncRNAs are significantly enriched in a specific molecular subtype of breast cancer (Fig. 1E). For example, SNHG15 is significantly enriched in the basal subtype (Fig. 1E and Fig. S2A-B). The regulation of SNHG7 and SNHG15 by IGF1 was confirmed with qPCR (Fig. 1F).

**Table 1.** SNHG7 is an lncRNA persistently regulated by IGF1 that is altered in breast cancer. Table indicates the expression, significant regulation (FDR <0.05) by IGF at 3 and 8hrs, alteration (copy number alterations and expression with z-Score threshold at +/- 2.0 in TCGA data as determined by cbioportal), and effect on survival (significant KM curve in altered vs. unaltered groups) of each persistently IGF-regulated lncRNA with a REFSeq ID.

**Hyper- Proliferation**
| IncRNA | Avg.FPKM | FC3hr | FC8hr | TCGA BC<br>%altered | TCGA BC<br>survival |
| --- | --- | --- | --- | --- | --- |
| **SNHG7** | 63.94 | -1.37 | -1.78 | 5 | yes |
| **SNHG1** | 49.98 | 1.63 | 1.69 | 8 | no |
| **SNHG15** | 17.62 | 2.75 | 3.10 | 7 | no |
| **HOTAIRM1** | 16.09 | -2.22 | 5.08 | 2 | no |
| **BCYRN1** | 5.85 | 1.95 | 2.33 | 2 | no |
| **LINC01138** | 5.56 | 2.11 | 1.94 | 0 | no |
| **MIR22HG** | 5.19 | 3.20 | 2.27 | 3 | no |
| **MIR210HG** | 3.30 | 2.35 | 3.09 | 1 | no |
| **LINC00324** | 2.35 | 2.35 | 2.06 | 2 | no |
| **SNHG12** | 11.18 | 1.57 | 1.64 | 4 | no |
| **MIR17HG** | 3.02 | 2.41 | 2.42 | 5 | no |
| **RPL23AP53** | 1.48 | 1.43 | 1.45 | 14 | no |

### SNHG7 is downregulated post transcriptionally by IGF via the MAPK pathway

Because SNHG7 is highly expressed, robustly regulated by IGF1 signaling, and is altered in a subset of breast cancer patients that correlate with survival, it was investigated further. SNHG7 is a relatively understudied lncRNA and is a <u>sn</u>oRNA <u>H</u>ost <u>G</u>ene (SNHG). SNHGs are highly structured genes (noncoding or protein coding) that have snoRNAs that are spliced and processed from their introns after they are transcribed, often resulting in two functional RNA species—1) snoRNAs and 2) mRNAs or lncRNAs. For example, the well-characterized tumor suppressor lncRNA GAS5, which is down-regulated in breast cancer, is a SNHG that has multiple snoRNAs that are processed from its introns. The snoRNAs are functional, but it is the mature GAS5 lncRNA that controls apoptosis by regulating glucocorticoid receptor signaling(24).

SNORA43 and SNORA17 are the snoRNAs expressed in two of the introns of SNHG7 (Fig. 2A and S3A). After the snoRNAs are spliced out of the primary SNHG7 transcript they are further processed to become functional snoRNAs. However, the mature SNHG7 transcript is conserved among primates (Fig. S3A), highly and ubiquitously expressed (Fig. S3B), unlikely to encode for a protein as indicated by low PhyloCSF(37) (negative for all 6 frames) and txCDsPredict (576.00) scores (both visualized in UCSC Genome Browser), and is predicted to be highly structured (Fig. S3C) suggesting it is noncoding and has biological functions independent of the snoRNAs. Both 5’ and 3’ Rapid Amplification of cDNA Ends (RACE) confirmed that there are at least two main REFseq annotated isoforms expressed in MCF7 cells that differ by one intron (Fig. 2A and Fig. S3A red and blue and S3D). In this report, the 5 exon, 4 intron isoform is referred to as SNHG7-I (Fig. S3A red) and the 4 exon, 3 intron isoform is referred to as SNHG7-NI (Fig. S3A blue). The 3^rd^ RefSeq SNHG7 isoform (Fig. S3A no color) was not detected by RACE. Subcellular fractionation followed by qPCR demonstrates SNHG7 is predominantly expressed in the cytoplasm (Fig. 2B).

**Figure 2.**
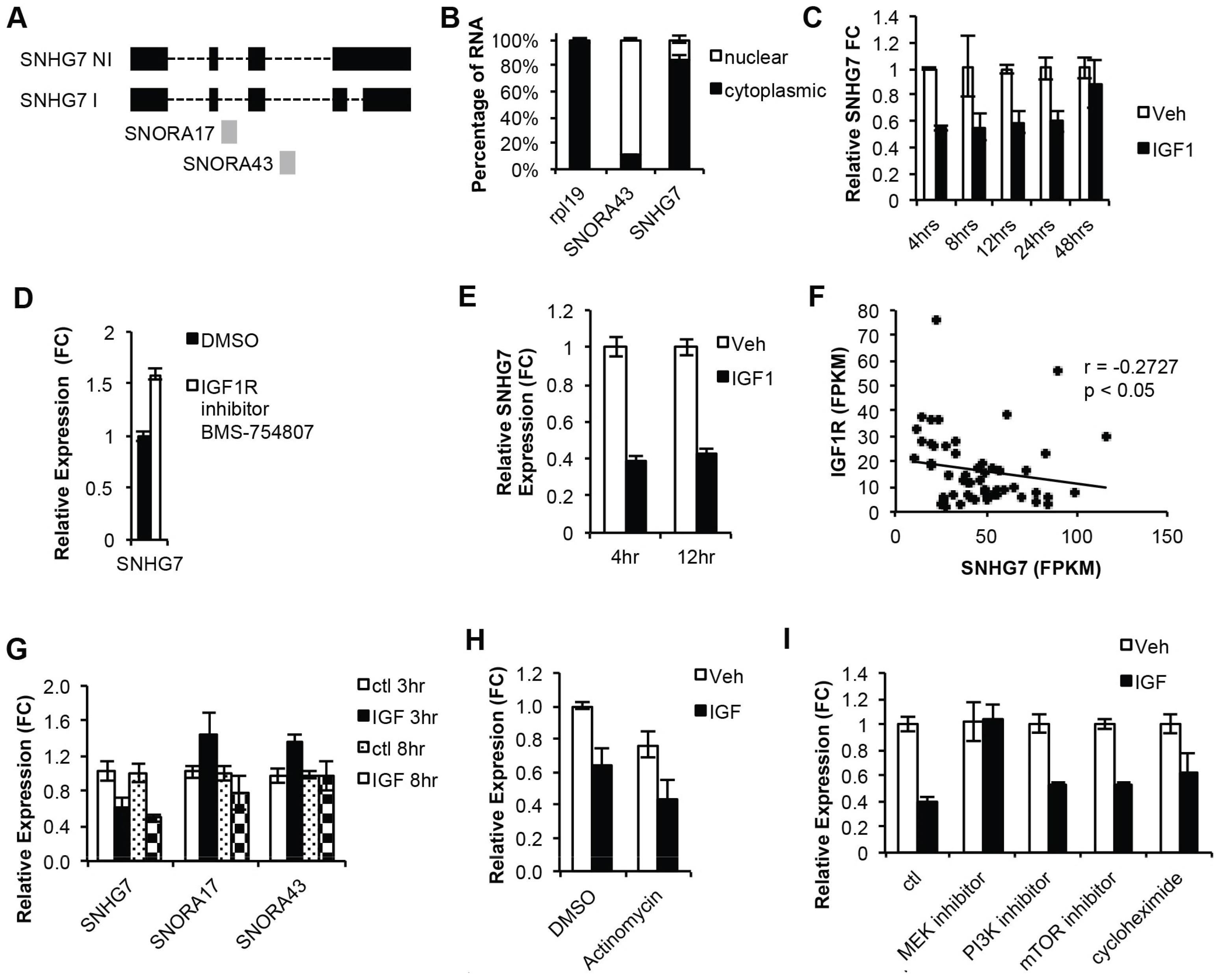
SNHG7 expression is downregulated by IGF1 signaling via a posttranscriptional mechanism through MAPK. (A) Schematic of two prominent isoforms of SNHG7 with (SNHG7 I) or without (SNHG7 NI) a fourth intron. SNORA17 and 43 are processed from the second and third introns of SNHG7. (B) RNA levels of the indicated genes in exponentially growing MCF7 cells following subcellular fractionation and subsequent qPCR analysis. The mean percentage +/- SD are reported. (C) Time course analysis of SNHG7 levels following the stimulation of serum starved MCF7 cells with 100nM IGF1 or vehicle control. Reported are the relative mean expressions +/- SD at each time point of biological triplicates to RPL19 and normalized to the respective vehicle treated cells. (D) MCF7 cells were plated in triplicate and the next day, BMS754807, an IGF1R inhibitor, was added to the media for 4 hours. RNA was isolated and qPCR performed as described above (t-test p<0.05). (E) SNHG7 levels determined by qPCR as in (C), but in MCF10a cells. (F) The correlation of expression of SNHG7 and IGF1R RNA in 56 breast cancer cell lines (spearman = −0.2727; p<0.05). Raw reads from RNAseq data published by Joe Grey et al. (38) from 56 breast cancer cell lines were reanalyzed through the pipeline described in Fig. 1A to determine the expression of SNHG7 and IGF1R. (G) Expression of the snoRNAs in SNHG7s introns determined by qPCR at 3 and 8 hrs. Levels were calculated as described above and are reported as the mean expression +/- SD of biological triplicates. (H-I) MCF7 cells were plated in triplicate for each treatment group, starved overnight, pretreated with the indicated drug for 1-2hrs before stimulation with IGF1 or vehicle control for 8hrs. Cells were harvested, RNA was isolated, cDNA was generated, and qPCR was performed and is presented as described above. (H) 10ug/ml of actinomycin was used to inhibit transcription and all results are normalized to the DMSO/Ctl group (I) 50uM of U0126 was used to inhibit MEK; 500nM of Wortmanin was used to inhibit PI3K; 1ug/ml of rapamycin was used to inhibit mTOR; 50 ug/ml of cycloheximide was used to inhibit translation; and, ctl was DMSO. Reported is the mean +/- SD normalized to the respective Ctl.

To test the kinetic regulation of SNHG7 by IGF1, MCF7 cells were treated with IGF1 for various lengths of time. The expression of SNHG7 is significantly and continuously down-regulated by IGF1 signaling for 24hrs (Fig. 2C). MCF7 cells were also treated with an IGF1R kinase inhibitor (BMS-754807) and the expression of SNHG7 increased, further implicating that the expression of SNHG7 is tightly regulated by IGF1 signaling (Fig. 2D). This regulation is not unique to MCF7 cells as SNHG7 is also regulated by IGF1 in the immortalized but non-transformed MCF10A cells (Fig. 2E). Additionally, there is a significant negative correlation (r=-0.2727;p<0.05) between RNA levels of SNHG7 and IGF1R (Fig. 2F) as determined by RNAseq data published for a set of 56 breast cancer cell lines(38) reanalyzed through the pipeline described above, suggesting the regulation of SNHG7 by IGF signaling is common in breast cancer cell lines.

While mature SNHG7 is downregulated by IGF1 signaling, the snoRNAs contained within the introns of SNHG7 are not significantly reduced (Fig. 2G), suggesting post-transcriptional regulation of mature SNHG7 instead of transcriptional regulation of the primary transcript. To determine if this is the case, serum starved MCF7 cells were treated with Actinomycin D before addition of IGF1 or vehicle. The inhibition of transcription did not ablate the reduction of SNHG7 expression by IGF1 (Fig. 2H) suggesting that IGF1 alters SNHG7 expression by reducing the stability of the transcript and not through transcriptional repression. The reduction of SNHG7 levels after Actinomycin treatment (Fig. 2H DMSO/Ctl vs. Actinomycin/Ctl) demonstrates transcription was effectively inhibited. Combined, these results suggest that the regulation of the mature transcript is not merely a mechanism to change the expression of the snoRNAs in the introns, but rather a tight regulation of the levels of the mature SNHG7 lncRNA.

SNHG7 is a 5’terminal oligopyrimidine (5’TOP) gene similar to Gas5. It is known that Gas5 lncRNA levels and other 5’TOP genes are destabilized by translation(39). Given that IGF1 signaling regulates translation, we tested if IGF1 regulates SNHG7 levels through translation. Surprisingly, we observed that inhibition of translation with cycloheximide did not prevent IGF1 from decreasing the levels of SNHG7 (Fig. 2I), so we examined the effects of signaling intermediates. Two of the primary downstream signaling pathways of IGF1R are PI3K/AKT/mTOR and MAPK. Small molecule inhibitors of PI3K, MEK, and mTOR were used to examine how IGF1 alters the stability of SNHG7. Inhibition of PI3K and mTOR had little effect on IGF1’s control of SNHG7 levels, while inhibition of MEK fully prevented alterations of SNHG7 levels by IGF1 signaling in serum starved MCF7 cells (Fig. 2I) indicating MEK signaling in the destabilization of SNHG7. Collectively, these results (Fig 2) suggest a novel mechanism whereby IGF1 significantly down-regulates the expression of SNHG7 through posttranscriptional alteration of SNHG7 mature RNA stability via the MAPK pathway.

### SNHG7 is necessary and sufficient for breast cancer cell proliferation

IGF1 signaling regulates proliferation of breast cancer cells. To determine if SNHG7 has similar effects, we examined the response of proliferation to altered SNHG7 levels. A pool of independently designed siRNA duplexes significantly reduced mature SNHG7 expression without altering the expression of the snoRNAs hosted in the introns (Fig. 3A). The proliferation of MCF7 cells with reduced SNHG7 expression was drastically reduced as scored by both a fluorometric assay measuring DNA content (Fig. 3B) and by counting cells with a hemacytometer using trypan blue exclusion (Fig. S4A-B). Proliferation of both other cell lines examined, MDA-MB-231 (Fig. S4C-D) and MCF10A (Fig. 3C) were also significantly reduced by RNAi targeting SNHG7. The inhibition of proliferation in these cells is due to the reduction of SNHG7 levels and not an off-target effect as demonstrated by the ability of 3 different individual siRNA duplexes (Fig. 3D) that target SNHG7 to all inhibit proliferation (Fig. 3E). Interestingly, these data suggest that there is a dose-dependent response to SNHG7 levels as the individual duplexes that were most efficient at inhibiting SNHG7 levels also inhibited proliferation the most (Fig. 3D-E). A live/dead assay demonstrated that the reduction in cell numbers by siSNHG7 treatment is due to a decrease in proliferation (Fig. S4E) and not an increase in cell death (Fig. S4F). While control treated cells continued to increase in number, siSNHG7 treated cells do not (Fig. S4E); however, the number of dead cells is not significantly different between treatment groups (Fig. S4F). Additionally, FACS analysis with propidium iodine staining indicates that by 3 days siSNHG7 treated MCF7 cells begin to arrest in G0/G1 (Fig. 3F). Reducing the expression of SNHG7 had no effect on the sensitivity of MCF7 cells to the dual-kinase IGF1R/InsR inhibitor, BMS-754807 (Fig.S4G). However, once again it is obvious that reduced SNHG7 expression decreases basal proliferation (Fig. S4G siCtl vs siSNHG7 at 10^-9^M). Together these data demonstrate that SNGH7 is necessary for full proliferation of breast cancer cell lines.

**Figure 3.**
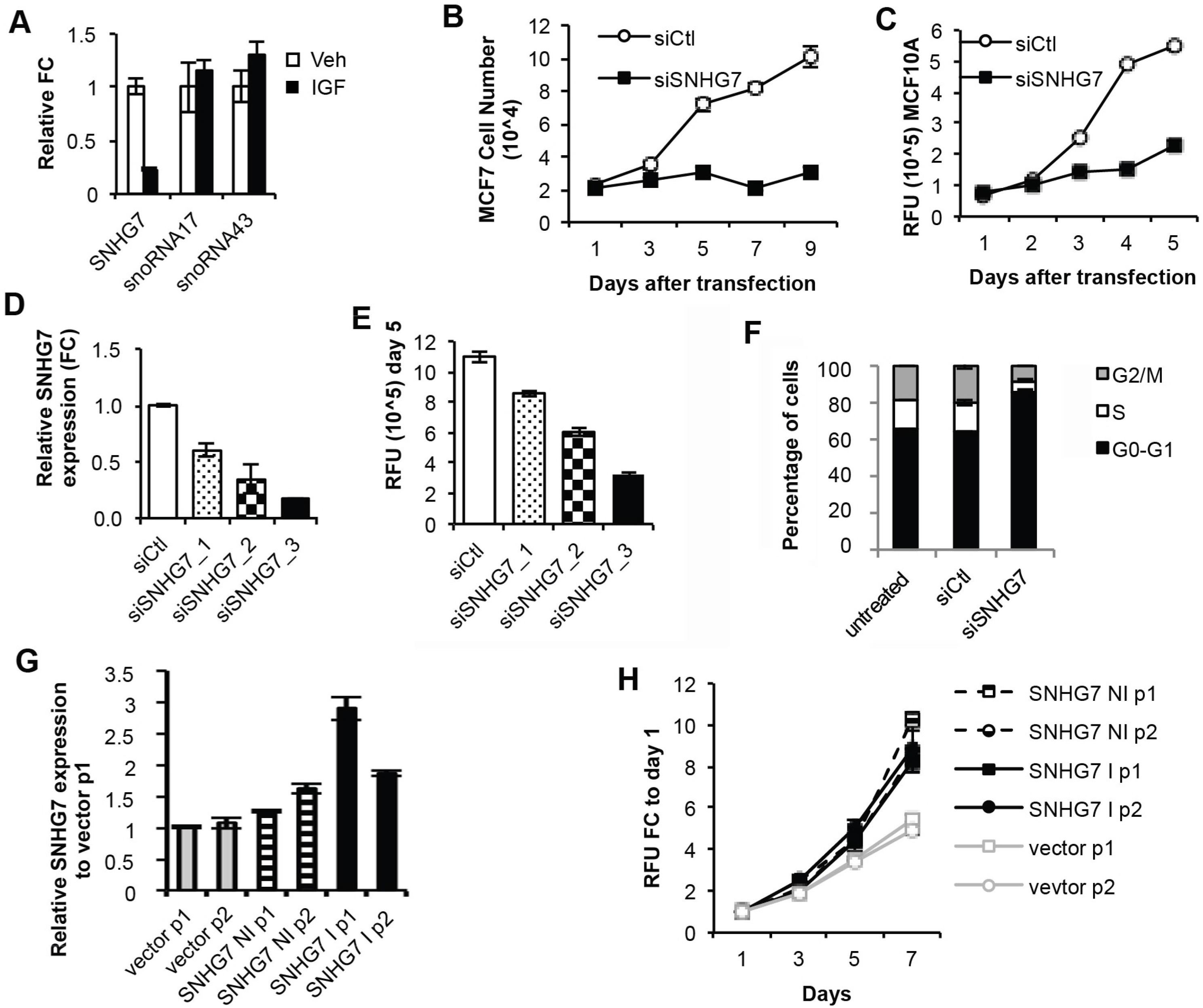
SNHG7 regulates proliferation in a dose-dependent manner. (A) MCF7 cells plated in triplicate were reverse transfected with a pool of two siRNA duplexes targeting SNHG7 or a non-targeting control. RNA was isolated and qPCR was performed as described earlier to determine expression levels of SNHG7 and the snoRNAs in its introns.(B-C) Eight biological replicates of (B) MCF7 or (C) MCF10a cells per treatment and time were reverse transfected as described above into 96-well dishes. At each time, media was removed and proliferation was assayed according to protocol (FluoReporter; Life) and mean +/- SEM is reported (non-linear regression; p<0.05). (D) MCF7 cells plated in triplicate were reverse transfected with three individual siRNA duplexes targeting SNHG7 or a non-targeting control (siCtl). RNA was isolated and qPCR was performed as described above to determine knockdown of SNHG7. (E) Eight biological replicates of MCF7 cells were reverse transfected with the three individual siRNAs for five days. Proliferation was measured as described above and the mean +/- SEM for 8 biological replicates are reported. All results in D and E are significant (ttests vs. siCtl <0.05). (F) MCF7 cells were reverse transfected in triplicate as described above. After 3 days the cells were fixed, stained with propidium iodide, and cell cycle analysis was performed using flow cytometry. The mean percentage of cells in each cell cycle phases +/- SD are graphed and are significantly different (ttest siCtl vs. siSNHG7; p<0.05). (G-H) The two isoforms of SNHG7 (see Fig. 2A) were cloned into pcdna3.1, transfected into MCF7 cells individually, and multiple polyclonal cell lines were generated by selection with G418. The number after p indicates the clone number. (G) qPCR was performed and mean +/- SEM are reported of biological triplicates to verify that SNHG7 was expressed higher than clones generated by transfection of vector alone (all significant; ttest p<0.05). (H) The proliferation of the MCF7 cells overexpressing either isoform of SNHG7 compared to empty vector was measured by the FluoReporter assay (normalized to day1 for each cell line to control for slight variation in seeding density). The mean +/- SEM is reported and shows that cells overexpressing either isoform of SNHG7 significantly (p<0.0001 nonlinear regression; 8 replicates for each treatment/time point) enhanced proliferation.

To test if SNHG7 is sufficient to induce or enhance proliferation, the two main isoforms of SNHG7 identified by RACE were cloned from cDNA of MCF7 cells. Two polyclonal MCF7 cell lines stably expressing SNHG7 were generated for each isoform (Fig. 3G) and non-linear regression analysis of proliferation data demonstrated that MCF7 cells overexpressing either isoform proliferated faster than cells expressing empty vector (Fig. 3H; doubling time=1.746-2.183 days for SNHG7 overexpressing cells vs. 2.684-2.89 days for empty vector cells p<0.0001). Therefore, SNHG7 is both necessary and sufficient for proliferation and regulates it in a dose-dependent manner. Furthermore, as described above, SNHG7 is overexpressed or amplified in ~5% of all breast cancer tumors in TCGA and correlates significantly with poorer disease-free survival (Fig. 1D and Table 1). This suggests that SNHG7 may act as an oncogene under certain conditions driving poor prognosis through the regulation of proliferation.

### IGF/SNHG7 feedback through regulation of common transcripts

Proliferation in response to IGF is regulated, at least in part, through the vast transcriptional changes downstream of IGF signaling. It is apparent that SNHG7 is also important for proliferation (Fig 3). To determine if SNHG7 controls proliferation through the alteration of similar transcripts as IGF1, we examined the expression of four known IGF1 regulated genes after knockdown of SNHG7 (versus scramble control) and in an SNHG7 overexpressing cell line (versus a vector control). Like IGF1 stimulation (Fig. 4A dark green), overexpression of SNHG7 (Fig. 4A dark blue) resulted in higher expression of LIF and EGR3 and lower expression of IRS2 and SOCS2 compared to empty vector control cells (Fig. 4A). Reduction of SNHG7 expression (Fig. 4A dark red) caused the opposite effect, decreased expression of LIF and EGR3 and increased expression of IRS2 and SOCS2 (Fig. 4A). Together these data suggest that IGF1 and SNHG7 regulate the expression and direction of expression of similar transcripts.

**Figure 4.**
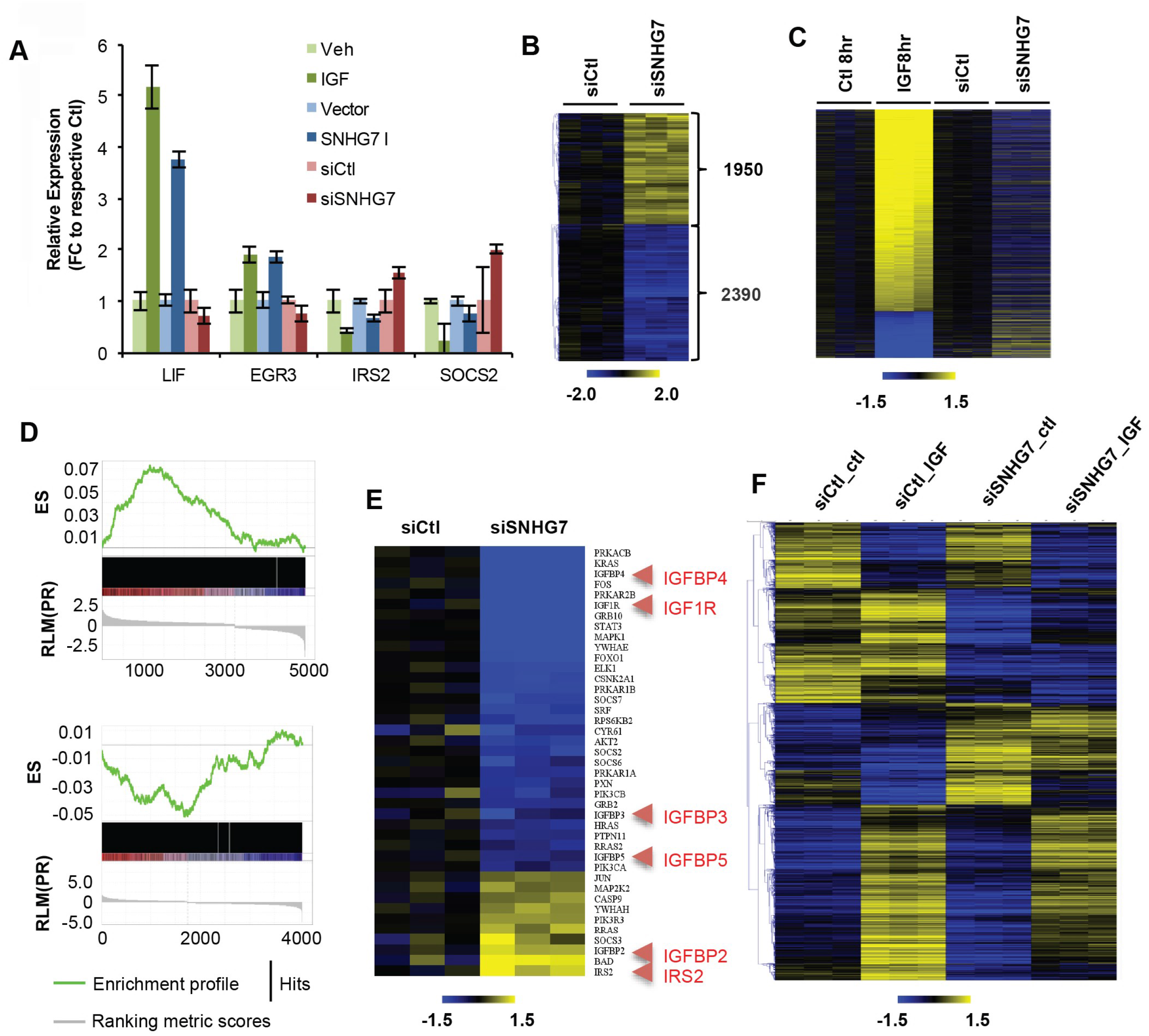
SNHG7 controls the expression of IGF1 signaling molecules and IGF1-regulated genes. (A) (green bars) MCF7 cells plated in triplicate were starved overnight and then treated with IGF1 or control. RNA was isolated, and qPCR was performed as described. The mean expression +/- SEM relative to control is reported to show example transcripts that are upregulated (Lif and Egr3) and downregulated (Irs2 and Socs2) by IGF1 signaling. (blue bars) The mean +/- SEM relative expression of the same genes from triplicate experiments in logarithmically growing MCF7 cells stably expressing SNHG7-I or a vector control to demonstrate regulation by overexpression of SNHG7. (red bars) Reverse transfection with siSNHG7 or control was performed as described previously and qPCR was performed to determine the expression of the same targets with decreased levels of SNHG7. (B) RNAseq was performed and analyzed as described in the methods following 3 days of siSNHG7 or siCtl treatment in MCF7 cells. The heatmap shows the relative expression of significantly regulated genes (q<0.05) for all replicates (N=3 for each condition) to the mean expression of Ctl treated cells. (C) Heatmap of significantly regulated genes by IGF after 8hrs of treatment of serum starved MCF7 cells and the respective expression of those genes following 3 days of siSNGH7 treatment. All expressions are normalized to the corresponding controls. (D) Gene Set Enrichment Preranked Analysis (GSEA) of (top) differentially expressed genes from IGF1RNAseq at 8hrs vs. differentially expressed genes from siSNHG7 RNAseq (FDR < 0.0001; Normalized Enrichment Score (NES) 2.83), and (bottom) GSEA of differentially expressed genes from siSNHG7 RNAseq vs. differentially expressed genes from IGF1 RNAseq at 8hrs (FDR < 0.05; NES = −1.84). ES = Enrichment Score. (E) Ingenuity Pathway Analysis revealed IGF1 signaling as a top pathway altered. Heatmap shows the 42 genes in the IGF1 pathway (out of 91) that are differentially regulated by siSNGH7 treatment. Highlighted red are key IGF1 signaling genes. (F) MCF7 cells reverse transfected for 2 days with siSNHG7 or nontargeting siCtl were serum starved overnight and treated with IGF or vehicle control (ctl) for 8hrs. RNA was isolated and RNAseq was performed as previously described. Shown are the log2 median centered values for all significantly altered genes (FDR < 0.05; average fpkm >1 for any condition; known annotation) between any of the conditions as determined by CuffDiff.

To examine if this pattern is comprehensive we performed RNAseq following reduced expression of SNHG7 by RNAi. The expressions of 4,341 genes were significantly altered (Fig. 4B and Supplementary Table 3; FDR <0.05) with 1308 annotated genes changing by at least 1.5-fold. The regulation of expression of several genes was confirmed with qPCR (Fig. S5). At a global level, there was a negative correlation between genes regulated by IGF1 induction and knockdown of SNHG7 (Fig. 4C). Gene Set Enrichment Analyses (GSEA) demonstrated that IGF1-regulated genes (8hrs; FDR<0.05) are highly enriched in genes regulated by siSNHG7 treatment (FDR <0.05; Fig. 4D top), and genes regulated by siSNHG7 are enriched for IGF1-regulated genes (Fig. 4D bottom). Collectively, these data demonstrate that IGF1 and SNHG7 control the transcript levels of a similar set of genes and suggest that SNHG7, in part, regulates proliferation through the control of a similar transcriptome response as IGF. Additionally, Ingenuity Pathway Analysis (IPA) of siSNHG7-regulated genes showed that the top canonical pathways are Molecular Mechanisms of Cancer and IGF1 Signaling (p=3.93E-09; 42/97 molecules altered; Fig. 4E) and the top molecular and cellular function is Cellular Growth and Proliferation. This further validates that SNHG7 is important in cancer development and proliferation. Likewise, it reveals that SNHG7 directly regulates the expression of IGF signaling transcripts (Fig. 4E) in addition to downstream targets in a manner that enhances the response of IGF1 signaling. However, RNAseq following IGF1 induction of siSNHG7 treated cells demonstrated that lack of SNHG7 did not prevent IGF from activating its signaling cascade (data not shown) or from regulating induction or repression of most transcripts (Fig. 4F; differences in siCtl_ctl and siCtl_IGF vs differences in siSNHG7_ctl vs. siSNHG7_IGF; Fig. S6). However, the overall levels of the transcripts were altered by reduction of SNHG7 expression leading to an attenuated IGF1 effect (Fig. 4F siCtl_ctl vs siSNGH7_ctl; Fig. 4F siCtl_IGF vs siSNHG7_IGF; Supplementary Table 4). This implies a fine-tuning feedback mechanism whereby IGF1 signaling decreases the expression of SNHG7, which is a positive regulator of IGF1 signaling intermediates and downstream targets through an independent regulation mechanism.

Finally, there are well-known issues with using breast cancer clinical data from TCGA due to short-term and limited follow up of the patients(40). Accordingly, we sought to confirm the clinical impact of extreme levels of SNHG7 in the tumors of breast cancer patients in the METABRIC(41) dataset that includes rich and long-term clinical data from over 2000 patients. However, the METABRIC gene expression dataset was calculated by microarray analysis, making it impossible to know the direct levels of SNHG7 and many other lncRNAs. For that reason, we used a guilt-by-association technique to infer the levels of SNHG7 in each of the patients. The top 100 upregulated and downregulated genes by siSNHG7, determined by fold change with a FDR <0.05, were used as an ‘SNHG7 signature’ and a Gene Set Variation Analysis(42) was performed to provide a score to each breast cancer tumor in the METABRIC dataset. Kaplan-Meier analysis demonstrates that patients with tumors with the highest decile of SNHG7 scores (indicative of high SNHG7 levels) have a significantly significant poorer disease-free survival (logrank test p-value=0.00079) than those with lower scores (Fig. S7). This further argues that SNHG7 has an important biological and clinical role in breast cancer.

## DISCUSSION

We leveraged the knowledge of IGF1 signaling and biology as a model system to identify a lncRNA, SNHG7, that is important for proliferation and breast cancer biology. By doing so we uncovered a novel fine-tuning feedback mechanism between IGF1 and SNHG7 that tightly regulates RNA expression and cell proliferation. As summarized in a schematic in Figure 5, our data shows that in addition to the regulation of many protein coding genes, IGF, which is necessary for proliferation, downregulates the expression of SNHG7. Our results also implicate SNHG7 in the regulation of expression of an enriched set of IGF1-regulated genes and of IGF1 signaling intermediates (Fig. 5 left). Additionally, there is a dose-response correlation between SNHG7 levels and proliferation. Therefore, when IGF1 signaling is active it alters gene expression (including downregulation of SNHG7) to increase proliferation (Fig. 5 middle). However, by reducing SNHG7, which regulates a similar set of genes as IGF1, and also numerous IGF1 signaling intermediates, the amplitude of IGF1-regulated genes is muted (Fig. 5 middle). When this feedback mechanism is overwhelmed, for example by the overexpression of SNHG7 or the disruption of SNHG7 regulation by IGF1 (indicated by an x), it leads to enhanced proliferation at least in part through differences in overall magnitude of IGF targets (Fig. 5 right – induced genes are expressed higher; repressed genes are repressed lower).

**Figure 5.**
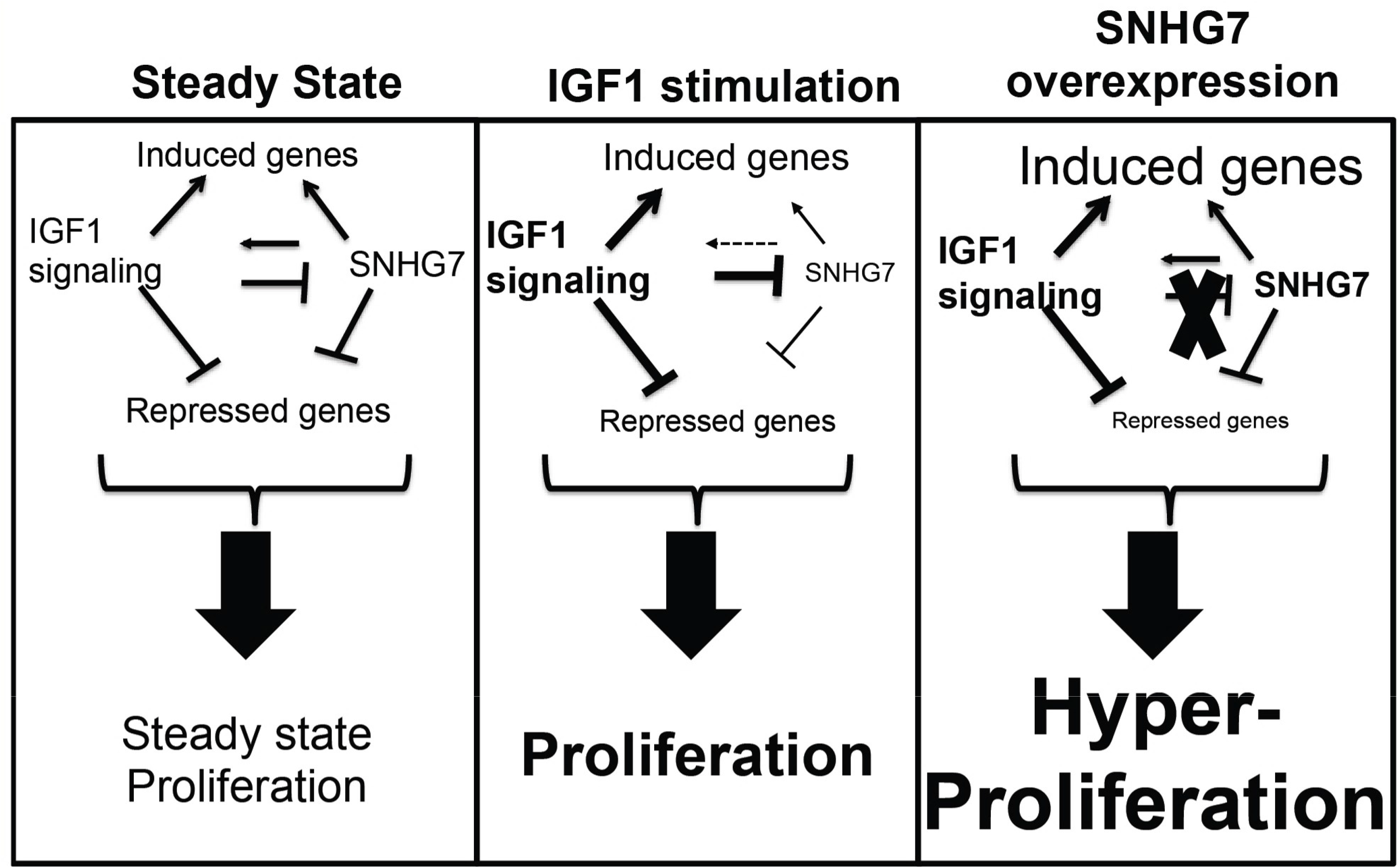
Model of attenuated regulation of IGF1 signaling and proliferation by SNHG7. (left) IGF signaling and SNHG7 regulate the expression of a similar gene set. IGF1 signaling decreases SNHG7 expression, while SNHG7 enhances the expression of IGF1 signaling molecules. Both IGF1 signaling and SNHG7 are necessary for proliferation. (middle) Upon enhanced IGF1 signaling, IGF1 initiates a transcriptional response, while simultaneously downregulating SNHG7, which attenuates the expression of the same transcriptional response; thus, a fine-tuning feedback mechanism that tightly regulates the proliferation response. (right) With overexpression of SNHG7 or the inability of IGF to downregulate SNHG7 as indicated by an X, the transcriptional response to IGF1 is enhanced (induced genes expressed higher; repressed genes expressed lower as indicated by the size of the font) leading to hyperproliferation.

It is paradoxical that IGF1 would repress SNHG7, which controls the expression of many of the same genes (in the same direction) and is necessary for proliferation, while simultaneously inducing proliferation. However, our results and others(9) show that IGF1 signaling reduces the expression of IRS2, an immediate downstream signaling scaffold, and increases the expression of numerous phosphatases (DUSPs) that dephosphorylate and inactivate many of the kinases downstream of IGF1R. Thus, IGF1 regulation of SNHG7 expression is an example of a systems biology feedback mechanism to auto-attenuate IGF1 signaling. Further, our knock-down experiments that completely inhibit proliferation reduce SNHG7 levels much lower than IGF1 signaling does (90% vs. 40%) suggesting there is a critical amount of SNHG7 necessary for proliferation. Therefore, we propose that IGF1 regulates SNHG7 levels as a feed-back mechanism to fine-tune the transcriptional response and proliferation induced by IGF1 to prevent hyperproliferation or transformation/progression. If this is true, we would predict that high levels of SNHG7 could lead to hyperproliferation. Accordingly, SNHG7 is overexpressed or amplified in ~5% of TCGA breast cancer patients, and these patients have worse disease-free survival than those without SNHG7 alterations.

In this report, we also describe a novel posttranscriptional mechanism of regulation of SNHG7 through alterations in stability via the MAPK pathway. SNHG7 is a 5’TOP gene like Gas5, which are regulated by nonsense mediated decay (NMD) through translation(43). While SNHG7 levels are altered by mTOR and translational inhibition (data not shown), it is clear that IGF1/MAPK regulation of SNHG7 levels is independent of translation induced by IGF1 because inhibition of translation, mTOR, and PI3K/AKT did not prevent IGF1 mediated downregulation of SNHG7. This suggests an additional mechanism of regulation of 5’TOP genes that requires further investigation.

Our results that IGF-regulated lncRNAs, including SNHG7 and SNHG15, are important for biology, enriched in breast cancer subtypes, and correlate with survival are consistent with recent studies. A large number of functionally important lncRNAs were shown to be regulated by estrogen signaling(25), but ours is the first study that examined regulation of lncRNAs by IGF. Additionally, through reanalysis of TCGA data, others have demonstrated that certain lncRNAs are enriched in specific breast cancer subtypes and lncRNAs alone can accurately stratify patients into molecular subtypes(44–46). In fact, lncRNAs were shown to be more subtype specific than protein coding genes and some correspond to patient survival, suggesting their utility as biomarkers(45). It is still unclear if SNHG7 or other IGF-regulated lncRNAs can be used as biomarkers or targeted for therapy. However, further understanding of the IGF1/SNHG7 system, the mechanisms of SNHG7 functions, and the characterization of other IGF1-regulated lncRNAs clearly will impact our understanding of both basic and breast cancer biology.

## Methods

### Cell Culture, treatments, and transfections

MCF7, MDA-MB-231, T47D, and MCF10A cells were obtained by ATCC and all experiments were performed within 25 passages. MCF7 and MDA-MB-231 cells were maintained in DMEM+10%FBS; T47D in RPMI-1640+10%FBS; and MCF10A cells in DMEM:F12(1:1)+5%HS, 20ng/ml EGF, 0.5 mg/ml hydrocortisone, 100ng/ml cholera toxin, and 10ug/ml Insulin. For IGF induction experiments, all cells were washed 2x in PBS and serum deprived in modified IMEM+10mM Hepes, 1ug/ml transferrin, 1ug/ml fibronectin, and 2mM l-glutamine for 16hrs before addition of 100ng/mL or equal volume of 10mM HCl as a vehicle control. To determine the mechanism of SNHG7 regulation, serum starved cells were pretreated for 1-2hrs with 10ug/ml actinomycin to prevent transcription, 1ug/ml rapamycin (mTOR inhibitor), 50uM U0126 (MEK inhibitor), 500 nM Wortmanin (PI3K inhibitor), or 50ug/ml cycloheximide (translational inhibitor) before addition of IGF1. BMS-754807 at 10 uM was used an IGF1R inhibitor. MCF7 cells expressing either SNHG7 isoform or vector alone were created by cloning and then transfecting (Fugene 6) the respective SNHG7 isoform from MCF7 generated cDNA using the GeneRacer Kit (Thermofisher) after 3’ RACE (see Supplementary Methods for primers and additional details). Individual polyclonal lines were isolated following 2 weeks of selection with 1ug/ml G418.

### RNA Sequencing

Total RNA from biological triplicates was isolated, quality was determined (Bioanalyzer), rRNA was depleted (RiboMinus), multiplexed paired-end libraries were prepared (Illumina TruSeq), and sequencing was performed on an Illumina HiSeq (IGF RNAseq) or NextSeq (siSNHG7 RNAseq). Quality of the sequencing was determined by running FastQC. Differential gene expression was calculated by mapping reads to hg19 with Tophat2 (masking reads to miRNAs, snRNAs, snoRNAs, rRNAs, and tRNAs) to a concatenated .gtf of UCSC known genes and lincRNA annotations published by the Broad Institute(48) and assembled using Cufflinks allowing for novel gene discovery. To determine IGF1 regulated lncRNAs as listed in Figure 1, the raw reads were reanalyzed to a newer and more comprehensive annotations. Reads were mapped to GRCh38 with Tophat2 as documented above using Gencode v.21 annotations and again assembled using Cufflinks allowing for novel gene discovery. For all analyses differential gene expression was determined with Cuffdiff and gene names were converted with custom scripts as needed. A conservative list of IGF-regulated lncRNAs was generated by extracting any differentially expressed gene with a Cufflinks prescribed lncRNA annotation (Gencode v.21). If that gene also had a protein coding gene annotation, it was not considered a lncRNA. Heatmaps of differentially expressed lncRNAs were generated in MeV after the described normalizations. Preranked Gene Set Enrichment Analysis (42) was performed according to instructions comparing IGF-regulated genes to those altered by siSNHG7 treatment. Ingenuity Pathway Analysis was performed according to protocol using genes significantly regulated (FDR <0.01) by siSNHG7 treatment compared to control. All reads are deposited in SRA with accession numbers: PRJNA514323, PRJNA515247, and PRJNA515028.

### Quantitative RT-PCR

After treatment at the indicated times, cells were harvested, RNA was isolated, cDNA was generated, and qPCR were performed as described previously(49). Relative RNA levels were calculated using the ΔΔCT method compared to RPL19 as the reference gene. All experiments were conducted in biological and technical triplicates. For subcellular localization, logarithmically growing cells were trypsinized, pellet was washed x2 in PBS, and cells were lysed in buffer RLN (50mM Tris-HCl pH 8.0, 140mM NaCl, 1.5 mM MgCl2, 0.5% NP40). After the cytoplasm was removed, the nuclear pellet was washed x2 in Buffer RLN before addition of buffer RLT (Qiagen). RNA from both fractions were isolated using Qiazol following manufactures’ protocol.

### RNA Interference

All cells were reverse transfected using 50-100 nM final concentration of either individual or 2-4 pooled oligos from Dharmacon (see Supplementary methods for sequences) using RNAi Max at a final concentration of 3ul/ml. All assays were performed ~72hrs after siRNA treatment.

### Proliferation Assays

Cells treated as described were seeded in 96-well dishes with at least 6 biological replicates. At the indicated times following treatment, plates were harvested and proliferation was scored with the FluoReporter (ThermoFisher) assay by quantitation of dsDNA according to manufacturers’ instructions on the Victor X4 (PerkinElmer). Proliferation was also scored via counting cells with a hemocytometer (Fig. S4A) using Trypan Blue exclusion in triplicate plated MCF7 cells in 6-well dishes.

### Cell Cycle Assay

MCF7 cells were reverse transfected with siSNHG7, nontargeting control, or nothing in biological triplicates. After 3 days, the cells were collected, fixed in 70% ethanol for 1hr, stained with 100ug/mL propidium iodide for 1hr, and then analyzed by flow cytometry. The percentage of cells in each phase of the cell cycle was calculated according to protocol.

## Supporting information

Supplemental Table 1. IGF RNAseq raw

Supplemental Table 2. IGF1 regulated lncRNA table 3 or 8 hrs

Supplemental Table 3. siSNHG7 vs siCtl rnaseq significant genes

Supplemental Table 4. siSNGH7 IGF rnaseq significant genes

Supplemental Table 5. qPCR primers

This article contains supporting information online at www.pnas.org

## ACKNOWLEDGEMENTS

We thank the Breast Cancer Research Foundation (AVL), National Cancer Institute of the National Institutes of Health award number R01CA94118 (AVL) and P30CA047904 (AVL), and a Susan G Komen for the Cure postdoctoral fellowship, award number PDF12229789 (DB). AVL is a recipient of Scientific Advisory Council award from Susan G. Komen for the Cure, and a Hillman Foundation Fellow. We would also like to thank William Horn and the Genomics Research Core at the University of Pittsburgh as well as McGill University for performing the RNA sequencing and Nolan Priedigkeit and Ryan Hartmaier for carefully editing the manuscript.

