## Supplemental Information for "A negative feedback loop between Insulin-like Growth Factor signaling and the lncRNA SNHG7 tightly regulates transcript levels and proliferation"

**This PDF file includes:**

Supplementary methods

Figs. S1 to S7

Captions for additional data tables S1 to S5

### Supplementary Methods

***RACE and cloning of SNHG7 isoforms*:** The 5' and 3' ends of SNHG7 were determined with 5' and 3' RACE using the GeneRacer Kit (Invitrogen) following manufacturers' instructions. Total RNA was isolated from logarithmically growing MCF7 cells. After ligation of GeneRacer RNA Oligo to decapped RNA, it was reverse transcribed using SuperScript III and GeneRacer Oligo dT Primer. Nested PCR according to protocol using the GeneRacer provided primers with the gene specific primers listed below was used to amplify 5' and 3' ends and then were resolved on agarose gel. The single band identified for 5' RACE and the two distinct bands for 3'RACE were cut, purified, and cloned using the TOPO TA Cloning kit for Sequencing. Multiple clones of each were sequenced to verify cDNA ends. The difference in the two 3'RACE bands was due to the presence of one intron as indicated in Fig. 2A and Fig. S3A and D. To clone the two full length SNHG7 isoforms, cDNA was generated as described above using GeneRacer Oligo dT Primer following ligation of GeneRacer RNA Oligo to decapped RNA. Nested PCR was performed with the primers listed below to generate full length SNHG7 with BamHI and NOTI restriction sites added at the 5' and 3' ends respectively. The PCR product was digested and ligated into pcDNA3.1(+) (Invitrogen) cut with the same restriction enzymes. The resulting plasmids were transformed and screened to obtain both main isoforms of SNHG7. Both isoforms were confirmed by sequencing.

***siRNA duplexes*:** all were purchased from Dharmacon with the On-Target-Plus modifications

siSNHG7 duplex 1 (5)- AGCCAGGAAGCUUCGGGAAUU
siSNHG7 duplex 2 (3)- CAACAGCCCUGAUCCAGAUUU
siSNHG7 duplex 3 (6)- AGAGUGACCAGGCUGACCAUU
siSNHG7 duplex 4 (7)- GAGGUGACUUCGCCUGUGAUU

The majority of experiments were performed using a pool of the most efficient siRNAs—siSNHG7 3 and 4 siSNHG15 – Lincode SMARTpool N-028841 (Dharmacon)

siCtl – On-Target-Plus Control Pool or pool of On-Target-Plus Control siRNA#1 and siRNA#2 as control against siSNHG7-3 and siSNHG7-4 pool used for most experiments.

***qPCR primers*:** see Supplementary Table 5.

***Cell Viability and Cytotoxicity*:** MCF7 cells were reverse transfected in replicates of 6 into 96-well dishes (one for each day) with siSNHG7 or non-targeting control. At the indicated times Viability/Cytotoxicity Reagent (ApoTox-Glo Triplex Assay;Promega) was added and fluorescence was measured according to protocol on the Victor X4 (PerkinElmer)

**Supplementary Figures**

**Supplementary Figure 1. RNAseq analysis demonstrates IGF1 regulates the expression of many RNAs.** (A) Venn diagram demonstrating the number of annotated genes (not novel) regulated (FDR <0.05; FC>1.5) by IGF1 treatment at (blue) 3 and (red) 8hrs. (B) Table showing number of genes regulated by IGF1 by different criteria. The numbers in parenthesis indicates the number of differentially expressed genes that are annotated in Gencode v.21. (C) Quantitative RT-PCR was performed on the identical RNA used for the RNAseq analysis and on an independent set of RNA harvested from similarly treated MCF7 cells. The mean fold changes at 3hrs (IGF/ctl) +/- SEM calculated by RNAseq and the two qPCR analyses are graphed for each mRNA tested. (D) Subset of Ingenuity Pathway Analysis of genes regulated at 3 and 8 hrs by IGF1. In the heatmap, blue indicates pathways downregulated (mostly Cell death), and orange indicates pathways upregulated.

**Supplementary Figure 2. SNHG15 is enriched in basal-like breast cancer.** (A) Chi-squared analysis on the number of basal-like tumors compared to all other molecular subtypes with or without overexpression/amplification of SNHG15 expression as determined by cBioPortal of TCGA Breast Cancer data ($p<0.0001$). (B) Box and whiskers plot of SNHG15 expression (normalized RSEM) of TCGA samples across molecular subtypes (one-way ANOVA; $p<0.05$).

**Supplementary Figure 3. SNHG7 is a primate conserved, broadly expressed, highly structured lncRNA with multiple isoforms. (A)** SNHG7 genomic structure in the UCSC genome browser shows the 3 RefSeq annotated isoforms of SNHG7. The two main isoforms identified by RACE (FigS3D) are indicated with colored boxes. The blue box shows the SNHG7-NI isoform that lacks the 4$^{th}$ intron. The red box indicates SNHG7-I that contains the 4$^{th}$ intron highlighted by the green box. The non-highlighted SNHG7 isoform was not detected by RACE. SNORA43 and SNORA17 are mature snoRNAs that are processed from the spliced introns of SNHG7. The bottom shows the conservation of SNHG7 and the snoRNAs across mammals. Blacklines indicate conservation. SNHG7 is highly conserved among primates. The snoRNAs and the 5’ promoter of SNHG7 are highly conserved across all mammals. (B) Expression of SNHG7 across tissues; downloaded from UCSC Genome Browser using GTEx data. (C) The predicted secondary structure of SNHG7 calculated by RNA Fold. The red indicates a strong prediction. (D) Agarose gel of nested-PCR results from 5’ RACE and 3’ RACE. The results were sequenced and the difference between the two 3’ RACE was the intron highlighted in A above.

**Supplementary Figure 4. SNHG7 is necessary for proliferation of breast cancer cell lines.** (A). Three replicates of MCF7 cells per treatment and time were reverse transfected. At each time, the number of floating and adherent cells were counted with a hemocytometer using Trypan Blue exclusion. The mean +/- SD of the total number of cells (living +dead) are recorded for each time and condition.  (B) Reduction of SNHG7 expression in A was confirmed with qPCR. (C) Six replicates of MDA-MB-231 cells for each time point were reverse transfected with siSNHG7 or nontargeting control. At the indicated times, media was removed and proliferation was assayed according to protocol (FluoReporter; Life) and mean +/- SEM is reported. (D)

Reduction of SNHG7 expression in MDA-MB-231 cells in C was confirmed with qPCR. (E-F) The number of living and dead MCF7 cells were determined at the indicated time points (6 replicates each) following reverse transfection of siRNA targeting SNHG7 or a non-targeting control according to protocol (ApoTox-Glo Triplex Assay;Promega). While siSNHG7 reduces the number of living cells (E) it is not cytotoxic (F). (G) Equally plated MCF7 cells were treated with siRNA targeting SNHG7 or control duplexes in 96-well dishes. The next day the indicated concentrations of BMS-754806 (IGF1R kinase inhibitor) were added to the cells (6 biological replicates). The mean +/- SEM cell number was determined using the CyQuant assay (ThermoFisher) 72hrs post treatment. The IC50 was not different between the conditions.

**Supplementary Figure 5. Validation of siSNHG7 RNAseq.** MCF7 cells plated in triplicate were treated with siSNHG7 or non-targeting control. After 72hrs, RNA was isolated, reverse transcribed, and quantified with qPCR for the indicated genes.

**Supplementary Figure 6. Reduced SNHG7 expression does not inhibit IGF1-induced gene expression.** RNAseq calculated expressions of genes altered by IGF1 stimulation in the presence or absence of siSNHG7 are shown normalized to the mean expression of their respective control (siCtl/IGF normalized to siCtl/Ctl and siSNHG7/IGF normalized to siSNHG7/Ctl). The vast majority of genes are induced or repressed by IGF regardless of siSNHG7 treatment, however the overall expression of the genes are significantly altered (Fig. 5F), which cannot be observed in this figure due to normalization to their respective controls.

**Supplementary Figure 7. High SNHG7 levels are significantly enriched in patients with worse Disease Free Survival.** The top 100 upregulated genes determined by fold change with a FDR <0.05 in the siSNHG7 RNAseq data were specified as 'downregulated genes' by SNHG7. Likewise, the top 100 downregulated genes were specified as 'upregulated genes.' The sign of the fold change was inverted with the assumption that overexpressing SNHG7 would have the opposite effect of reducing SNHG7 expression. These 200 genes were defined as the SNHG7 signature and used to calculate ssGSEA scores for all 1980 tumors in the METABRIC dataset according to protocol. KM plots were generated in R with the top decile of SNHG7 scores vs the bottom 90% of tumors. The logRankP=0.00079 and N at each main time point is indicated in the figure.

**Supplementary Table 1**.

RNAseq data following IGF1 treatment of serum starved MCF7 cells and analysis through the Tuxedo Package.

**Supplementary Table 2.**

Table of IGF1 regulated lncRNAs at either 3 or 8 hours after IGF1 treatment in MCF7 cells.

**Supplementary Table 3.**

RNAseq data of siSNGH7 vs siCtl treatment in MCF7 cells. Analysis is through the Tuxedo package as described in the Methods Section.

**Supplementary Table 4.**

RNAseq data of IGF1 and vehicle treatment in siSNGH7 and siCTL treated MCF7 cells. Analysis is through the Tuxedo package as described in the Methods Section.

**Supplementary Table 5.**

List of qPCR primers.
