## Supplemental Figures for "A negative feedback loop between Insulin-like Growth Factor signaling and the lncRNA SNHG7 tightly regulates transcript levels and proliferation"

A

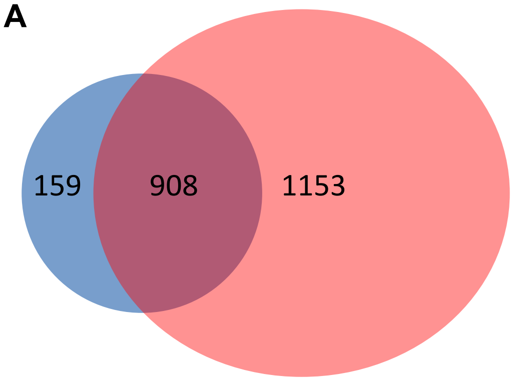


B

| Criteria | 3hr | 8hr |
|---|---|---|
| q<0.05 | 2593 (2076) | 4206 (3692) |
| q<0.05;FC>=1.5 | 1151 (1067) | 2492 (2061) |
| q<0.05;FC>=2.0 | 638 (353) | 881 (617) |

C

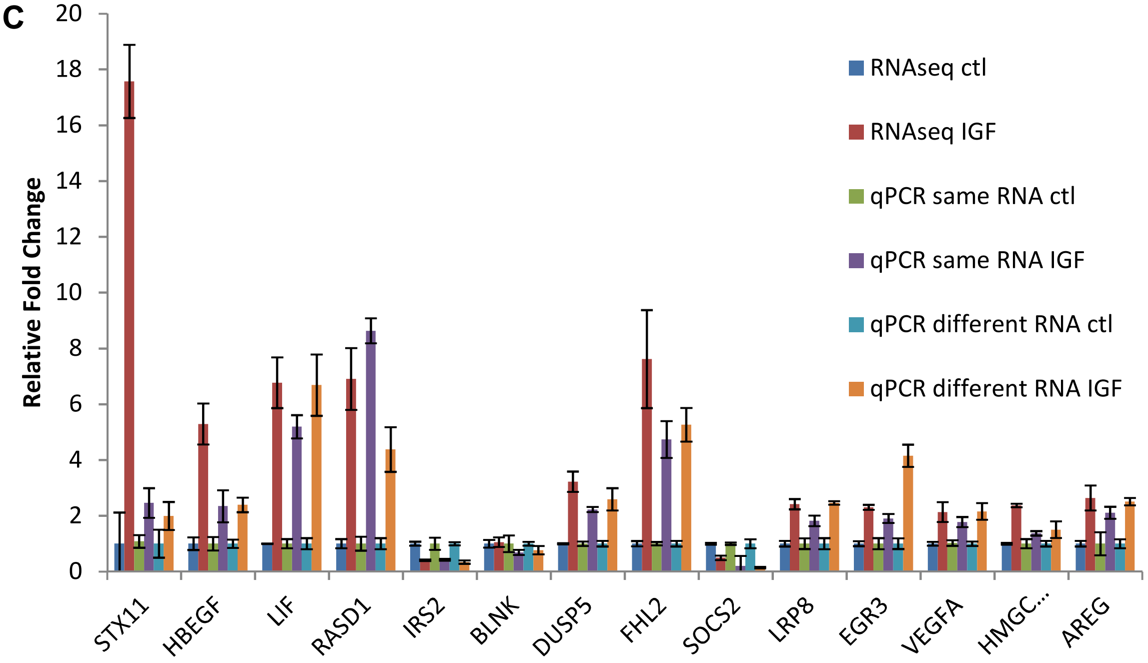


D

IPA of genes regulated at 3 and 8 hrs

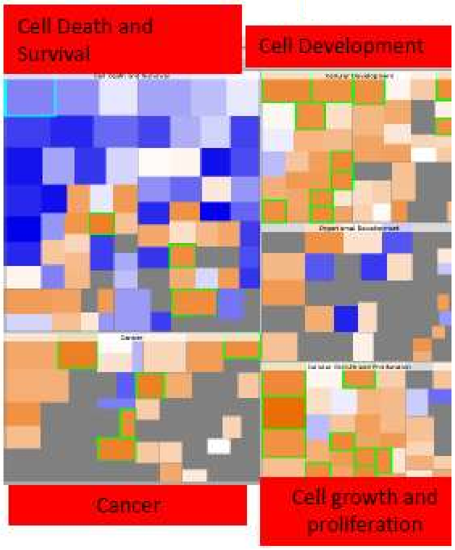


| ID | Associated Network Functions | Score |
|---|---|---|
| 1 | Cell Cycle, RNA Post-Transcriptional Modification, Cell Death and Survival | 45 |
| 2 | Cancer, Reproductive System Disease, Post-Translational Modification | 38 |
| 3 | Cellular Development, Cancer, Cell Death and Survival | 38 |
| 4 | Glomerular Injury, Renal and Urological Disease, Tissue Development | 34 |
| 5 | Cellular Development, Cellular Growth and Proliferation, Connective Tissue Development and Function | 33 |

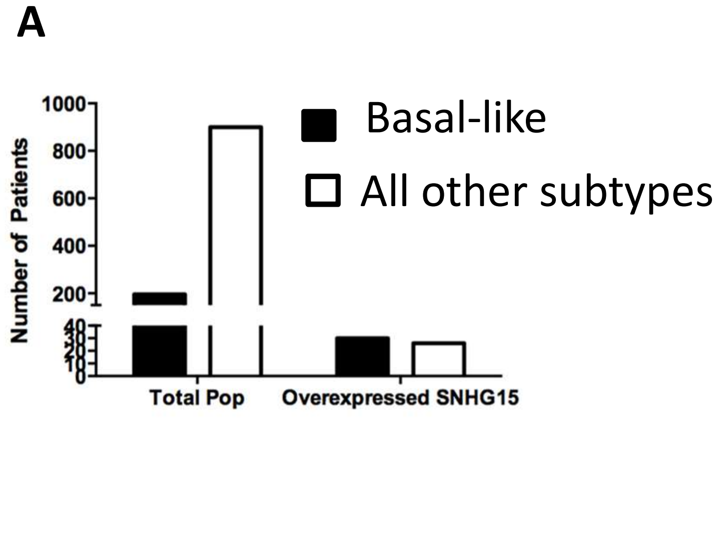
A
B
Number of Patients
1000
800
600
400
200
40
30
20
10
0
Total Pop
Overexpressed SNHG15
Basal-like
All other subtypes
SNHG15
*
Expression
1500
1000
500
0
Basal-like
Her2
LumA
LumB
Normal-like
Subtype

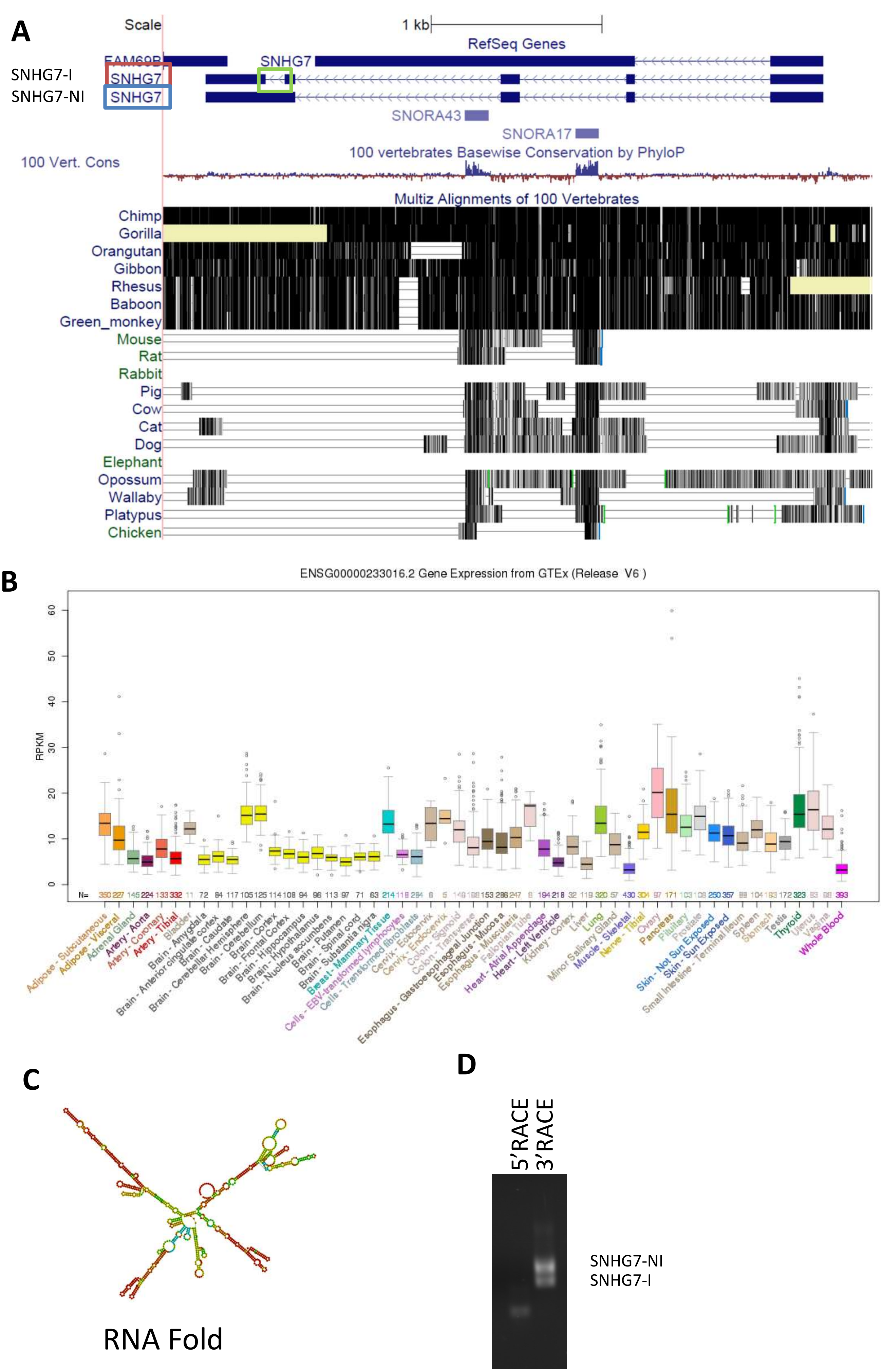
A
Scale
1 kb
RefSeq Genes
FAM69B
SNHG7
SNHG7-I
SNHG7
SNHG7-NI
SNHG7
SNORA43
SNORA17
100 Vert. Cons
100 vertebrates Basewise Conservation by PhyloP
Multiz Alignments of 100 Vertebrates
Chimp
Gorilla
Orangutan
Gibbon
Rhesus
Baboon
Green_monkey
Mouse
Rat
Rabbit
Pig
Cow
Cat
Dog
Elephant
Opossum
Wallaby
Platypus
Chicken
B
ENSG00000233016.2 Gene Expression from GTEx (Release V6 )
RPKM
0
10
20
30
40
50
60
N=
Adipose - Subcutaneous
Adipose - Visceral
Adrenal Gland
Artery - Aorta
Artery - Coronary
Artery - Tibial
Bladder
Brain - Amygdala
Brain - Anterior cingulate cortex
Brain - Caudate
Brain - Cerebellar Hemisphere
Brain - Cerebellum
Brain - Cortex
Brain - Frontal Cortex
Brain - Hippocampus
Brain - Hypothalamus
Brain - Nucleus accumbens
Brain - Putamen
Brain - Spinal cord
Brain - Substantia nigra
Breast - Mammary Tissue
Cells - EBV-transformed lymphocytes
Cells - Transformed fibroblasts
Cervix - Ectocervix
Cervix - Endocervix
Colon - Sigmoid
Colon - Transverse
Esophagus - Gastroesophageal Junction
Esophagus - Mucosa
Esophagus - Muscularis
Fallopian Tube
Heart - Atrial Appendage
Heart - Left Ventricle
Kidney - Cortex
Liver
Lung
Minor Salivary Gland
Muscle - Skeletal
Nerve - Tibial
Ovary
Pancreas
Pituitary
Prostate
Skin - Not Sun Exposed
Skin - Sun Exposed
Small Intestine - Terminal Ileum
Spleen
Stomach
Testis
Thyroid
Uterus
Vagina
Whole Blood
C
RNA Fold
D
5'RACE
3'RACE
SNHG7-NI
SNHG7-I

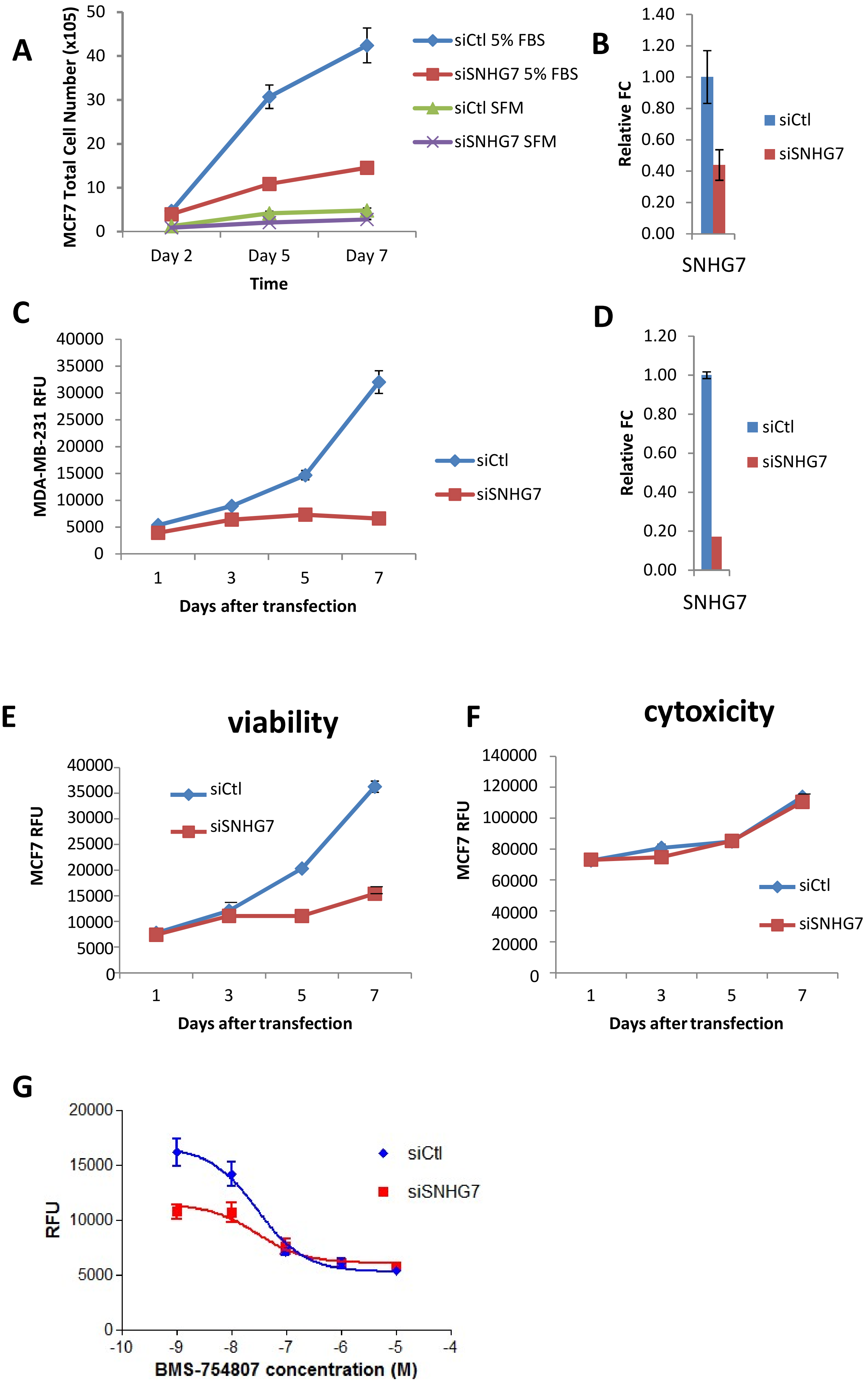
A
MCF7 Total Cell Number (x105)
50
40
30
20
10
0
Day 2
Day 5
Day 7
Time
siCtl 5% FBS
siSNHG7 5% FBS
siCtl SFM
siSNHG7 SFM
B
Relative FC
1.40
1.20
1.00
0.80
0.60
0.40
0.20
0.00
SNHG7
siCtl
siSNHG7
C
MDA-MB-231 RFU
40000
35000
30000
25000
20000
15000
10000
5000
0
1
3
5
7
Days after transfection
siCtl
siSNHG7
D
Relative FC
1.20
1.00
0.80
0.60
0.40
0.20
0.00
SNHG7
siCtl
siSNHG7
E
viability
MCF7 RFU
40000
35000
30000
25000
20000
15000
10000
5000
0
1
3
5
7
Days after transfection
siCtl
siSNHG7
F
cytoxicity
MCF7 RFU
140000
120000
100000
80000
60000
40000
20000
0
1
3
5
7
Days after transfection
siCtl
siSNHG7
G
RFU
20000
15000
10000
5000
0
-10
-9
-8
-7
-6
-5
-4
BMS-754807 concentration (M)
siCtl
siSNHG7

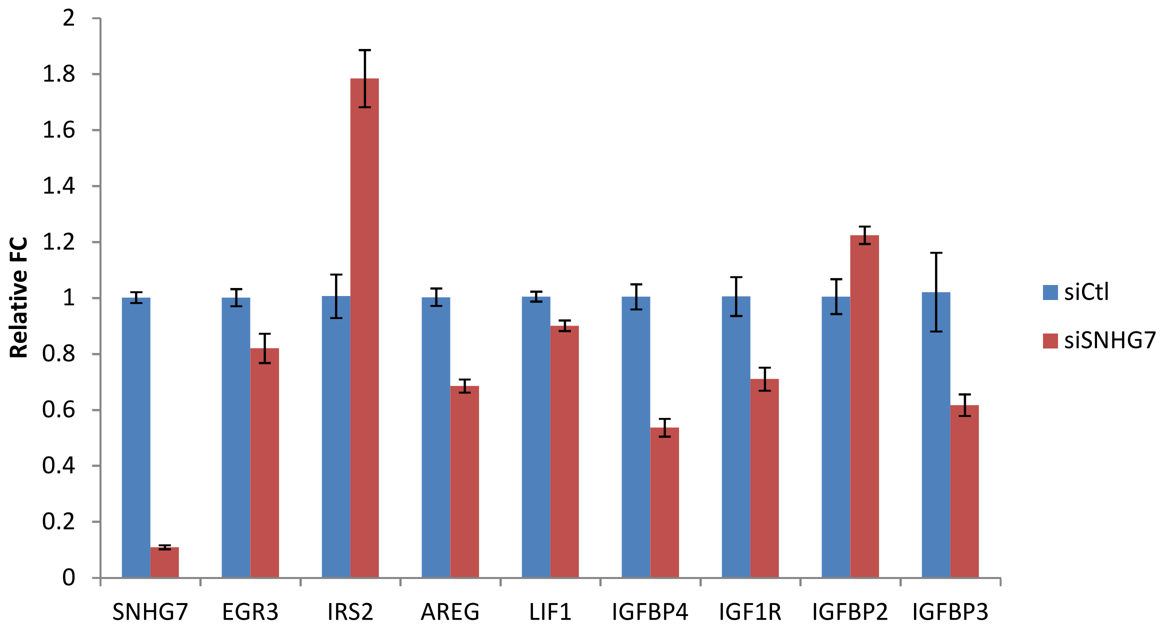
Relative FC
2
1.8
1.6
1.4
1.2
1
0.8
0.6
0.4
0.2
0
SNHG7
EGR3
IRS2
AREG
LIF1
IGFBP4
IGF1R
IGFBP2
IGFBP3
siCtl
siSNHG7

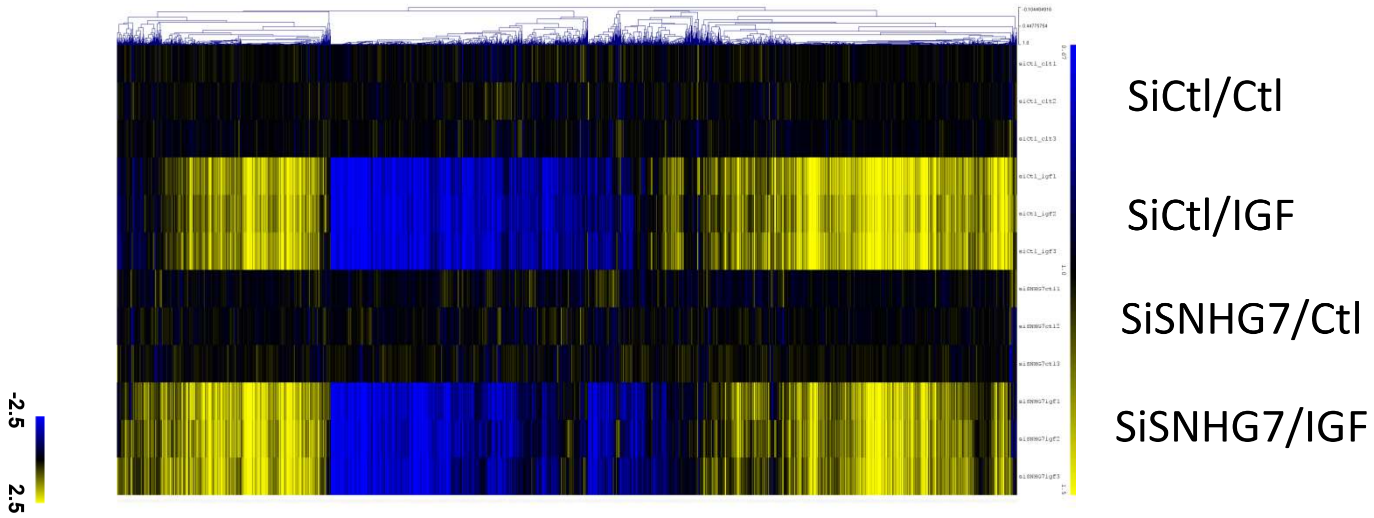
SiCtl/Ctl
SiCtl/IGF
SiSNHG7/Ctl
SiSNHG7/IGF
-2.5
2.5

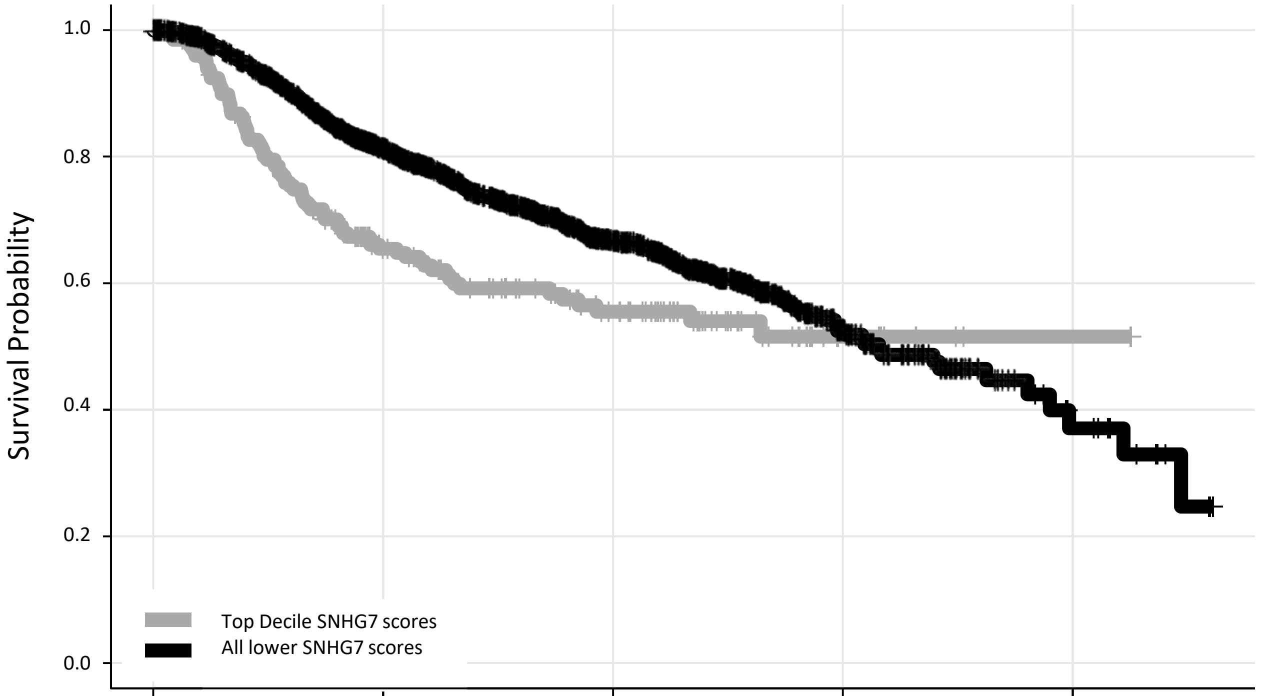


N at risk

| | | | | | |
|---|---|---|---|---|---|
| High | 198 | 101 | 54 | 9 | 1 |
| Low | 1782 | 1134 | 533 | 84 | 13 |
